## Supplemental Model S4 for "Cell Wall Dynamics in a Filamentous Fungus"

### Derivation of governing equations

We describe the dynamics of the exocytic vesicle level, *EV*, strain rate at tip, *G*, and thickness at tip, *h*. The surface growth speed is equal to the strain rate at the tip multiplied by the square of radius of curvature at tip, *R*. We also use as an intermediate variable the concentration in the cell wall (at the tip) of remodelling factors, *c*. We aim at deriving a parsimonious model for the dynamics of the growing tip.

Strain rate is proportional to the elastic strain *PR*/*Yh* in excess of a threshold $\epsilon$, known as yield strain, (Lockhart, 1965) and to the concentration of remodelling factors (Rojas and Dumais, 2011), with a coefficient $\mu$:

$$\boxed{G = \mu c \left(\frac{PR}{Yh} - \epsilon\right)_+}. \qquad (0)$$

The + subscript indicates the positive part of what precedes. In the following, we drop the subscript, but for numerical simulations, we set G'(t)=0 if G<0 or $\frac{PR}{Yh} - \epsilon$<0, to ensure growth occurs only if there is excess strain.

The rate of feeding of the exocytic vesicles is sensitive to the strain rate (feedback parameter $\phi$) and the exocytic vesicle level decays by transfer to the cell wall (rate $\alpha$):

$$\boxed{\frac{dEV}{dt} = \phi G - \alpha EV}. \qquad (1)$$

Remodelling factors are incorporated in the cell wall from the exocytic vesicles (incorporation coefficient $\beta$) and are diluted by growth (Rojas and Dumais, 2011). The concentration of cell remodelling factors per unit surface, *h c,* is then described by

$$\boxed{\frac{d(hc)}{dt} = \beta EV - Ghc}. \qquad (2')$$

The thickness increases primarily by synthesis of polysaccharides at the membrane, with number of synthases assumed to be proportional to the exocytic vesicle level, and secondarily by incorporation of polysaccharides synthesized in the Golgi and transiting through exocytic vesicles, at rate proportional to the level of exocytic vesicle; altogether, the thickness increases at a rate proportional to *EV*, with a coefficient $\gamma$. The thickness is diluted by growth (Rojas and Dumais, 2011; Davi et al. 2018). Therefore,

$$\boxed{\frac{dh}{dt} = \gamma EV - Gh}. \qquad (3)$$

We finally rewrite (2') in terms of strain rate, by using (0) to substitute *c* and (3) to substitute d*h*/d*t*. We obtain

$$\boxed{\frac{dG}{dt} = \eta EV \frac{1-\theta h}{h^2} - \gamma \frac{2-\theta h}{(1-\theta h)h} G EV + \frac{1}{1-\theta h} G^2}, \qquad (2)$$

in which the new parameters are

$$\boxed{\begin{aligned} \eta &= \frac{\beta \mu PR}{Y} \\ \theta &= \frac{\epsilon Y}{PR} \end{aligned}}. \qquad (4)$$

We note that the condition that wall expansion occurs (positive excess strain) amounts to $h\,\theta < 1$ .

In view of the osmotic shocks performed below, we note that, from Eq. (0), a discontinuity $\Delta P$ in pressure, results in a discontinuity of strain rate, given by

$$\boxed{\Delta G = \frac{G}{1-\theta h}\,\frac{\Delta P}{P}} \qquad (5)$$

Equations (1), (2), and (3) define a system of ordinary differential equations of order 1 for exocytic vesicle level, *EV*, strain rate, *G*, and tip thickness, *h*. These equations have 5 positive parameters: mechanical feedback coefficient $\phi$, decay rate of exocytic vesicle level $\alpha$, effective remodelling rate $\eta$, wall synthesis coefficient $\gamma$, and effective threshold $\theta$.

### Stationary points and their stability

We first use the right hand side of Eqs. (1-3) to write the function that defines the dynamical system.

```
In[1]:= DynSys[EV_, G_, h_] :=
         {ϕ G - α EV, η EV (1 - θ h) / h^2 - γ (2 - θ h) / (1 - θ h) / h G EV + G^2 / (1 - θ h),  γ EV - G h}
```

We seek the stationary points.

```
In[4]:= StationaryPoints = Solve[DynSys[EV, G, h] == 0, {EV, G, h}]
```

Solve: Equations may not give solutions for all "solve" variables.

Out[4]= $\left\{\{EV \to 0, G \to 0\}, \left\{EV \to 0, G \to 0, h \to \frac{\gamma\,\phi}{\alpha}\right\}, \left\{EV \to \frac{\eta\,(\alpha - \gamma\,\theta\,\phi)}{\alpha\,\gamma^2}, G \to -\frac{\eta\,(-\alpha + \gamma\,\theta\,\phi)}{\gamma^2\,\phi}, h \to \frac{\gamma\,\phi}{\alpha}\right\}\right\}$

All points with zero *EV* and *G* are stationary (no-growth stationary points). There is a single non-degenerate stationary point (with growth).

We then study their stability, by computing the Jacobian.

```
In[5]:= Jac[EV_, G_, h_] :=
         Transpose[{D[DynSys[s, g, H], s], D[DynSys[s, g, H], g], D[DynSys[s, g, H], H]}] /.
          {s → EV, g → G, H → h}
```

We first consider the stability of the stationary points with no growth.

```
In[6]:= JacNoGrowth = Jac[EV, G, h] /. StationaryPoints[[1]] // Simplify
        Eigenvalues[JacNoGrowth] // Simplify
```

Out[6]= $\left\{\{-\alpha, \phi, 0\}, \left\{\frac{\eta - h\,\eta\,\theta}{h^2}, 0, 0\right\}, \{\gamma, -h, 0\}\right\}$

Out[7]= $\left\{0, -\frac{h\,\alpha + \sqrt{h^2\,\alpha^2 + 4\,\eta\,\phi - 4\,h\,\eta\,\theta\,\phi}}{2\,h}, \frac{-h\,\alpha + \sqrt{h^2\,\alpha^2 + 4\,\eta\,\phi - 4\,h\,\eta\,\theta\,\phi}}{2\,h}\right\}$

This point is unstable because $h\,\theta < 1$, hence the third eigenvalue is positive.
This means that growth may start if the level of exocytic vesicle becomes strictly positive, which corresponds to the initiation of polarization.

We then consider the stability of the stationary point with growth.

In[8]:=
```
JacNoGrowth = Jac[EV, G, h] /. StationaryPoints[[3]] // Simplify
Eigenvalues[JacNoGrowth] // Simplify
```

Out[8]= $\left\{\{-\alpha, \phi, 0\}, \left\{-\frac{\alpha^2 \eta}{\gamma^2 \phi^2}, \frac{\eta \theta}{\gamma}, 0\right\}, \left\{\gamma, -\frac{\gamma \phi}{\alpha}, \frac{\eta (-\alpha + \gamma \theta \phi)}{\gamma^2 \phi}\right\}\right\}$

Out[9]= $\left\{\frac{\eta (-\alpha + \gamma \theta \phi)}{\gamma^2 \phi}, -\frac{\alpha \gamma - \eta \theta + \frac{\sqrt{2 \alpha \gamma \eta \theta \phi + \eta^2 \theta^2 \phi + \alpha^2 (-4 \eta + \gamma^2 \phi)}}{\sqrt{\phi}}}{2 \gamma}, \frac{1}{2}\left(-\alpha + \frac{\eta \theta + \frac{\sqrt{2 \alpha \gamma \eta \theta \phi + \eta^2 \theta^2 \phi + \alpha^2 (-4 \eta + \gamma^2 \phi)}}{\sqrt{\phi}}}{\gamma}\right)\right\}$

In[10]:= $2 \alpha \gamma \eta \theta + \eta^2 \theta^2 + \alpha^2 \left(\gamma^2 - \frac{4 \eta}{\phi}\right) - (-\alpha \gamma + \eta \theta)$ `^2 // Simplify`

Out[10]= $-\frac{4 \alpha \eta (\alpha - \gamma \theta \phi)}{\phi}$

The first eigenvalue is negative because $h\,\theta < 1$ and $h = \frac{\gamma \phi}{\alpha}$; the second and third eigenvalues are negative if $\theta < \alpha \gamma / \eta$.
Accordingly, **this stationary point is stable as long as $\theta$ is small enough** (this condition is not very restrictive because we already assume $\theta < 1/h$).

We thus keep working with this stationary solution. We note that the first eigenvalue is the same as strain rate. The ratio between the third eigenvalue (which is larger than the second) and the first can be written in the form $\frac{1}{2x(1-y)}\left(1 - x y - \sqrt{1 - 4x + 2xy + x^2 y^2}\right)$, which is either not real (so entails oscillations) or is larger than one. As no overshoot is observed in experimental data, this suggests that the first eigenvalue (equal to strain rate) controls the transient dynamics of the system.

In[11]:=
```
EVSta = EV /. StationaryPoints[[3]]
GSta = G /. StationaryPoints[[3]]
hSta = h /. StationaryPoints[[3]]
EigenvaluesNoGrowth = Eigenvalues[JacNoGrowth] //
  Simplify[#, Assumptions → {α > 0, γ > 0, ϕ > 0, η > 0, θ > 0}] &
```

Out[11]= $\frac{\eta\,(\alpha-\gamma\,\theta\,\phi)}{\alpha\,\gamma^2}$

Out[12]= $-\frac{\eta\,(-\alpha+\gamma\,\theta\,\phi)}{\gamma^2\,\phi}$

Out[13]= $\frac{\gamma\,\phi}{\alpha}$

Out[14]= $\left\{\frac{\eta\,(-\alpha+\gamma\,\theta\,\phi)}{\gamma^2\,\phi},\ -\frac{\alpha\,\gamma-\eta\,\theta+\sqrt{2\,\alpha\,\gamma\,\eta\,\theta+\eta^2\,\theta^2+\alpha^2\left(\gamma^2-\frac{4\,\eta}{\phi}\right)}}{2\,\gamma},\ \frac{-\alpha\,\gamma+\eta\,\theta+\sqrt{2\,\alpha\,\gamma\,\eta\,\theta+\eta^2\,\theta^2+\alpha^2\left(\gamma^2-\frac{4\,\eta}{\phi}\right)}}{2\,\gamma}\right\}$

### Estimation of parameters for hyphae in reference conditions

We have 5 parameters and so we need 5 equations to determine them. The value of the stationary states for *EV*, *G*, and *h* provide 3 equations. We then use an equation that is related to the growth threshold, provided by the arrest of growth post osmotic treatment. Given that no overshoot is observed in the experimental data, the time scale of transients is given by strain rate and so does not provide additional information, we therefore take as a first guess the limit between over-damped and underdamped oscillations, i.e. assume the second and third eigenvalue to be equal, and then we study the sensitivity of the results to this choice.

We use $\mu$m and min for units of length and time.

1) We take:
Stationary exocytic vesicle level $EV = 1$,
Stationary growth rate $G = Gs/R$ =0.6 $\mu$m/min/0.86$\mu$m=0.69 min $^{-1}$,
Stationary tip thickness $h = 0.066$ $\mu$m,
Relative drop in pressure that arrests growth: p=14%. From Eqs. (1) and (4), $\theta = \frac{1-p}{h}$ = 13.03$\mu$m $^{-1}$
We then obtain values of $\gamma$, $\eta$ and a relation between $\alpha$ and $\phi$.

2) We assume the two last eigenvalues to be equal and obtain the value of $\phi$, as justified hereafter.

In[19]:=
```
ConstraintsFromData = {EVData → 1, GData → 0.69, hData → 0.066, θ → 13.03}
Solve[{EVSta == EVData, GSta == GData, hSta == hData}, {γ, η, α}] // Simplify
ParametersFromData = % /. ConstraintsFromData // Flatten
EigenvaluesWithParameters =
 EigenvaluesNoGrowth /. ParametersFromData /. ConstraintsFromData // Simplify
```

Out[19]= {EVData → 1, GData → 0.69, hData → 0.066, θ → 13.03}

Out[20]= $\left\{\left\{\gamma \to \frac{\text{GData hData}}{\text{EVData}}, \eta \to \frac{\text{GData}^2\,\text{hData}^2}{\text{EVData} - \text{EVData hData}\,\theta}, \alpha \to \frac{\text{GData}\,\phi}{\text{EVData}}\right\}\right\}$

Out[21]= $\{\gamma \to 0.04554, \eta \to 0.0148114, \alpha \to 0.69\,\phi\}$

Out[22]= $\left\{-0.69, 2.11893 - 0.345\,\phi - 10.9794\sqrt{0.0372461 - 0.0160782\,\phi + 0.00098738\,\phi^2},\right.$
$\left. 2.11893 - 0.345\,\phi + 10.9794\sqrt{0.0372461 - 0.0160782\,\phi + 0.00098738\,\phi^2}\right\}$

In[23]:=
```
Plot[Re[EigenvaluesWithParameters] // Evaluate,
 {ϕ, 0, 20}, PlotLegends → {"v1", "Re(v2)", "Re(v3)"}]
```

Out[23]=
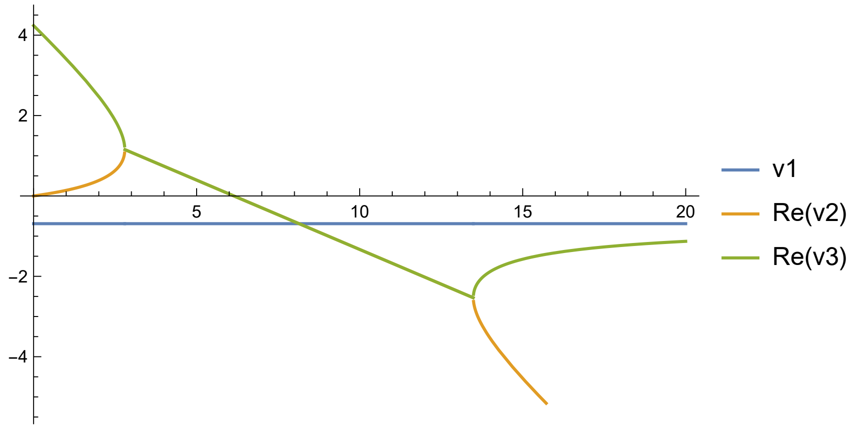


The first eigenvalue of the Jacobian is constant and equal to -*G*. We then consider the second and third ones. We observe three regimes for these two eigenvalues in the plot above: real and positive for small $\phi$ (the condition $\theta<\alpha\,\gamma/\eta$ is not satisfied); complex conjugates for intermediate $\phi$; real and smaller than *G* for large $\phi$. Based on the guess above, we use the value, $\phi_g$, of $\phi$ at the transition between the second and third regime. The value of $\phi$ has no influence on the equilibrium points and it is expected that, within the third regime, it has limited influence on the dynamics because the slowest mode is the eigenmode associated with the first eigenvalue. Indeed, for all results shown hereafter, we explored the range $\phi_g$ to 5 $\phi_g$ and found no qualitative change and relatively small quantitative changes.

In[24]:=
```
Solve[EigenvaluesWithParameters[[2]] == EigenvaluesWithParameters[[3]], ϕ]
ϕguess = %[[2, 1]]
```

Out[24]= {{ϕ → 2.79699}, {ϕ → 13.4867}}

Out[25]= ϕ → 13.4867

### Model with no feedback

Before going further, we examine whether a model with no feedback is compatible with experimental data. We replace Eq. (1) by

$$\boxed{\frac{dEV}{dt} = \phi - \alpha\, EV}. \qquad (1')$$

in which we assumed constant feeding of the exocytic vesicles, instead of mechanosensitive feeding.

We use Eqs. (1', 2, 3) to write the function that defines the dynamical system.

```
In[26]:= DynSysNoFeedback[EV_, G_, h_] :=
  {ϕ - α EV, η EV (1 - θ h) / h^2 - γ (2 - θ h) / (1 - θ h) / h G EV + G^2 / (1 - θ h), γ EV - G h}
```

We seek the stationary points.

```
In[27]:= StationaryPointsNoFeedback = Solve[DynSysNoFeedback[EV, G, h] == 0, {EV, G, h}]
```

Out[27]= $\left\{\left\{EV \to \frac{\phi}{\alpha},\ G \to -\frac{\gamma\,\eta\,\theta\,\phi}{-\alpha\,\eta + \gamma^2\,\phi},\ h \to \frac{\alpha\,\eta - \gamma^2\,\phi}{\alpha\,\eta\,\theta}\right\}\right\}$

There is a single stationary point as long as $\alpha\,\eta - \gamma^2\,\phi > 0$ and $\theta$>0; there is no stationary point otherwise.

We then study its stability, by computing the Jacobian.

```
In[28]:= JacNoFeedback[EV_, G_, h_] :=
  Transpose[{D[DynSysNoFeedback[s, g, H], s], D[DynSysNoFeedback[s, g, H], g],
     D[DynSysNoFeedback[s, g, H], H]}] /. {s → EV, g → G, H → h}
JacNoFeedbackSationaryPoint =
  JacNoFeedback[EV, G, h] /. StationaryPointsNoFeedback[[1]] // Simplify
Eigenvalues[JacNoFeedbackSationaryPoint] // Simplify
```

Out[29]= $\left\{\{-\alpha, 0, 0\},\ \left\{-\frac{\alpha^2\,\eta^3\,\theta^2}{(\alpha\,\eta - \gamma^2\,\phi)^2},\ \frac{\eta\,\theta}{\gamma},\ 0\right\},\ \left\{\gamma,\ \frac{-\alpha\,\eta + \gamma^2\,\phi}{\alpha\,\eta\,\theta},\ \frac{\gamma\,\eta\,\theta\,\phi}{-\alpha\,\eta + \gamma^2\,\phi}\right\}\right\}$

Out[30]= $\left\{\frac{\gamma\,\eta\,\theta\,\phi}{-\alpha\,\eta + \gamma^2\,\phi},\ -\alpha,\ \frac{\eta\,\theta}{\gamma}\right\}$

The third eigenvalue is positive and so the stationary point is unstable (if it exists). This model does not allow stable stationary growth and this would also apply to any modification of Eq. (1') in which the right-hand side is solely a function of *EV*.

### Estimation of parameters of the model with no feedback

The situation is the same as in the model with feedback, except that we do not use the argument on oscillations and $\phi$ remains as a free parameter.

```
In[31]:= EVStaNoFeedback = EV /. StationaryPointsNoFeedback[[1]]
GStaNoFeedback = G /. StationaryPointsNoFeedback[[1]]
hStaNoFeedback = h /. StationaryPointsNoFeedback[[1]]
Solve[{EVStaNoFeedback == EVData, GStaNoFeedback == GData, hStaNoFeedback == hData},
  {γ, η, α}] // Simplify
ParametersFromDataNoFeedback = % /. ConstraintsFromData // Flatten
```

Out[31]= $\frac{\phi}{\alpha}$

Out[32]= $-\frac{\gamma\,\eta\,\theta\,\phi}{-\alpha\,\eta+\gamma^2\,\phi}$

Out[33]= $\frac{\alpha\,\eta-\gamma^2\,\phi}{\alpha\,\eta\,\theta}$

Out[34]= $\left\{\left\{\gamma\to\frac{\text{GData hData}}{\text{EVData}},\ \eta\to\frac{\text{GData}^2\ \text{hData}^2}{\text{EVData}-\text{EVData hData}\ \theta},\ \alpha\to\frac{\phi}{\text{EVData}}\right\}\right\}$

Out[35]= {γ → 0.04554, η → 0.0148114, α → ϕ}

### Simulation of the model with no feedback

We illustrate the case of no feedback by making a simulation with a chosen value of $\phi$ (e.g. 5). If we add a small perturbation to the stationary solution (e.g. increasing or decreasing the initial value of *EV* by 0.001), then either thickness vanishes (cell wall rupture) or growth stops (with thickening of the wall).

```
In[36]:= ϕchoice = ϕ → 5
sol =
 First@NDSolve[{EV'[t] == ϕ - α EV[t], G'[t] == If[G[t] > 0, η EV[t] (1 - θ h[t]) / h[t]^2 -
          γ (2 - θ h[t]) / (1 - θ h[t]) / h[t] G[t] × EV[t] + G[t]^2 / (1 - θ h[t]), 0],
       h'[t] == γ EV[t] - G[t] × h[t] , EV[0] == EVData - 0.0001, G[0] == GData,
       h[0] == hData} /. ConstraintsFromData /.
     ParametersFromDataNoFeedback /. ϕchoice, {EV, G, h}, {t, 0, 20}]
Plot[EV[t] /. sol, {t, 0, 2.78}, PlotRange → All, AxesLabel → {"t(min)", "EV(AU)"}]
Plot[G[t] /. sol, {t, 0, 2.78}, PlotRange → All, AxesLabel → {"t(min)", "G(1/min)"}]
Plot[10^3 h[t] /. sol, {t, 0, 2.78}, PlotRange → All, AxesLabel → {"t(min)", "h(nm)"}]
```

Out[36]= ϕ → 5

NDSolve: At t == 2.7852855053556986`, step size is effectively zero; singularity or stiff system suspected.

Out[37]= {EV → InterpolatingFunction[ Domain: {{0., 2.79}} Output: scalar ],

G → InterpolatingFunction[ Domain: {{0., 2.79}} Output: scalar ],

h → InterpolatingFunction[ Domain: {{0., 2.79}} Output: scalar ]}

Out[38]=

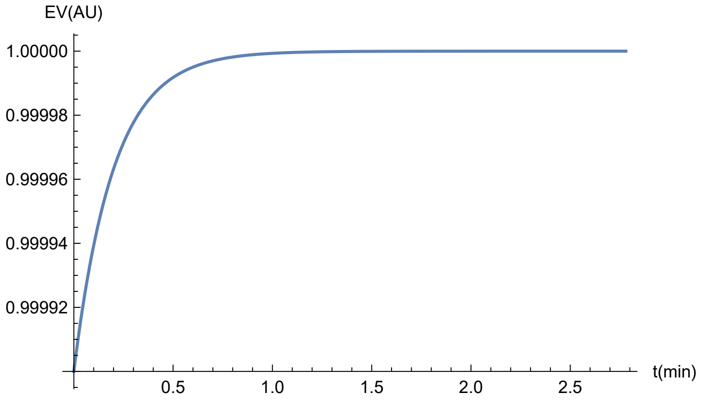


Out[39]=

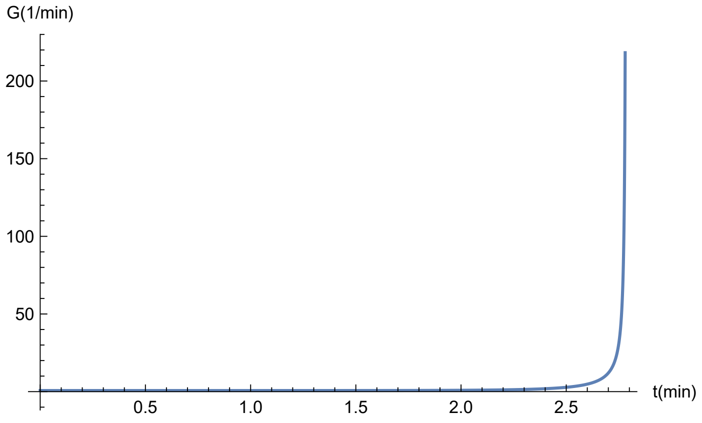


Out[40]=

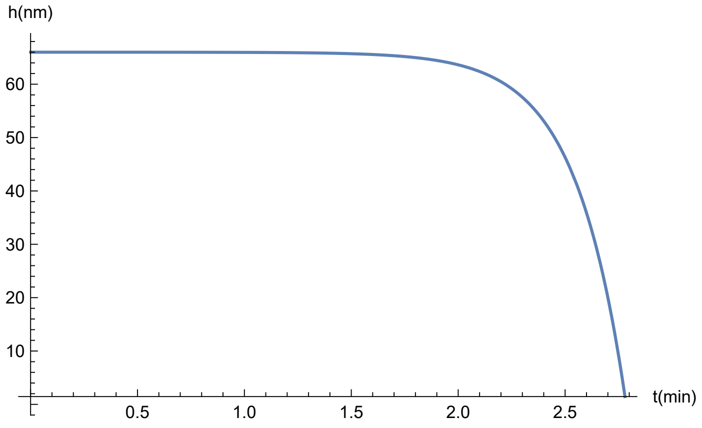

```
Export[
 "/Users/arezki/Documents/Recherche/Yeast/Louis/SimulationGraphs/NoFeedback1.xlsx",
 Table[{t, EV[t] /. sol, G[t] /. sol, 10^3 h[t] /. sol}, {t, 0, 2.78, 0.01}]]
```

Out[•]= /Users/arezki/Documents/Recherche/Yeast/Louis/SimulationGraphs/NoFeedback1.xlsx

In[41]:=
```
ϕchoice = ϕ → 5
sol =
 First@NDSolve[{EV'[t] == ϕ - α EV[t], G'[t] == If[G[t] > 0, η EV[t] (1 - θ h[t]) / h[t]^2 -
          γ (2 - θ h[t]) / (1 - θ h[t]) / h[t] G[t] × EV[t] + G[t]^2 / (1 - θ h[t]), 0],
      h'[t] == γ EV[t] - G[t] × h[t] , EV[0] == EVData + 0.00001,
      G[0] == GData, h[0] == hData} /. ConstraintsFromData /.
    ParametersFromDataNoFeedback /. ϕchoice, {EV, G, h}, {t, 0, 20}]
Plot[EV[t] /. sol, {t, 0, 20}, PlotRange → All, AxesLabel → {"t(min)", "EV(AU)"}]
Plot[G[t] /. sol, {t, 0, 20}, PlotRange → All, AxesLabel → {"t(min)", "G(1/min)"}]
Plot[10^3 h[t] /. sol, {t, 0, 20}, PlotRange → All, AxesLabel → {"t(min)", "h(nm)"}]
```

Out[41]= ϕ → 5

Out[42]= {EV → InterpolatingFunction[ Domain: {{0., 20.}} Output: scalar ],
G → InterpolatingFunction[ Domain: {{0., 20.}} Output: scalar ],
h → InterpolatingFunction[ Domain: {{0., 20.}} Output: scalar ]}

Out[43]=
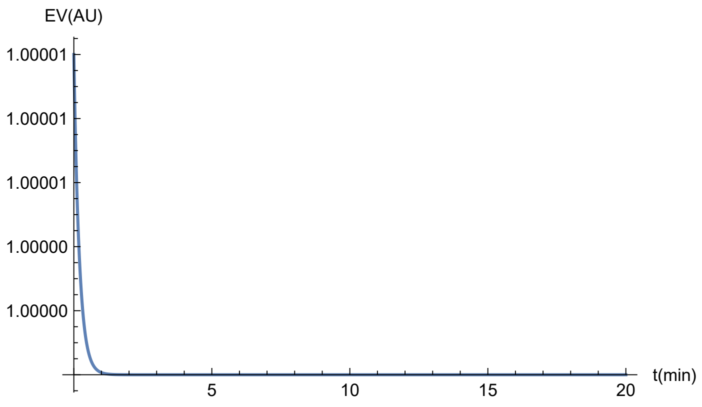


Out[44]=
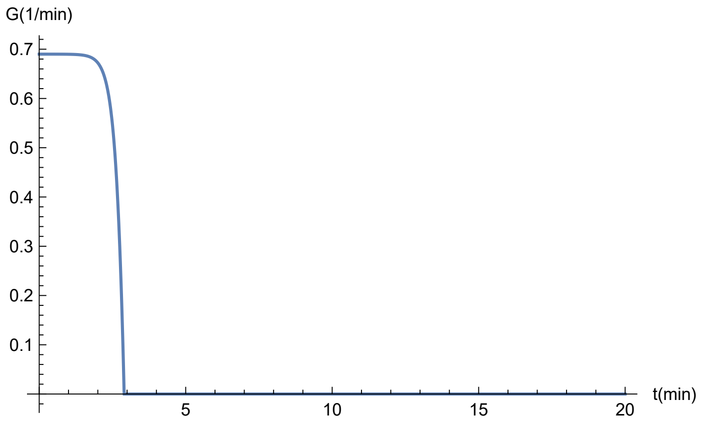

Out[45]=

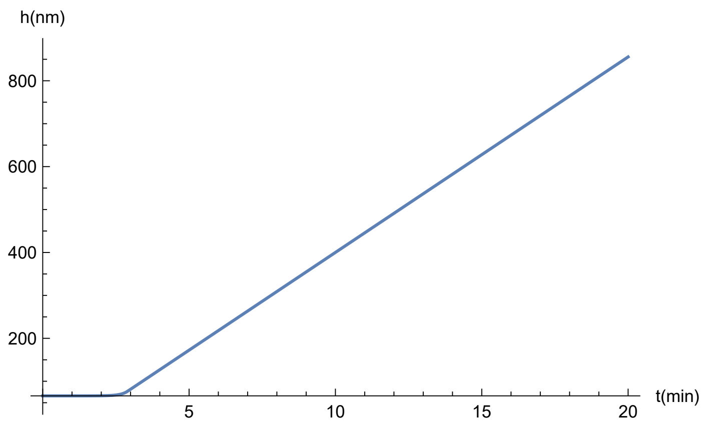


```
Export[
 "/Users/arezki/Documents/Recherche/Yeast/Louis/SimulationGraphs/NoFeedback2.xlsx",
 Table[{t, EV[t] /. sol, G[t] /. sol, 10^3 h[t] /. sol}, {t, 0, 20, 0.01}]]
```

Out[∘]= /Users/arezki/Documents/Recherche/Yeast/Louis/SimulationGraphs/NoFeedback2.xlsx

```
Export["D:\\_Stage_Minc_Levure\\Model\\SimulationGraphs/Nofeeback1_Rtip.xlsx",
 Table[{t, EV[t] /. sol, G[t] /. sol, 10^3 h[t] /. sol}, {t, 0, 20, 0.01}]]
```

Out[∘]= D:_Stage_Minc_Levure\Model\SimulationGraphs/Nofeeback1_Rtip.xlsx

### Simulation of side branching

We start with initiation of growth. We solve numerically the dynamical system, starting with low exocytic vesicle level, no growth, and reference value of thickness. We find that polarity and growth are established within about 10-15 minutes, a delay comparable to that in experiments.
*Note*: the units of plots are min for time, arbitrary for *EV*, min $^{-1}$ for strain rate, and nm for thickness.

In[46]:=
```
sol = First@
   NDSolve[{EV'[t] == ϕ G[t] - α EV[t], G'[t] == If[G[t] > 0, η EV[t] (1 - θ h[t]) / h[t]^2 -
             γ (2 - θ h[t]) / (1 - θ h[t]) / h[t] G[t] × EV[t] + G[t]^2 / (1 - θ h[t]), 0],
        h'[t] == γ EV[t] - G[t] × h[t] , EV[0] == .1, G[0] == 0, h[0] == hData} /.
      ConstraintsFromData /. ParametersFromData /. ϕguess, {EV, G, h}, {t, 0, 20}]
Plot[EV[t] /. sol, {t, 0, 20}, PlotRange → All, AxesLabel → {"t(min)", "EV(AU)"}]
Plot[G[t] /. sol, {t, 0, 20}, PlotRange → All, AxesLabel → {"t(min)", "G(1/min)"}]
Plot[10^3 h[t] /. sol, {t, 0, 20},
 PlotRange → {0, 1.1 × 10^3 hData /. ConstraintsFromData},
 AxesLabel → {"t(min)", "h(nm)"}]
```

Out[46]= {EV → InterpolatingFunction[ Domain: {{0., 20.}} Output: scalar ],

G → InterpolatingFunction[ Domain: {{0., 20.}} Output: scalar ],

h → InterpolatingFunction[ Domain: {{0., 20.}} Output: scalar ]}

Out[47]=

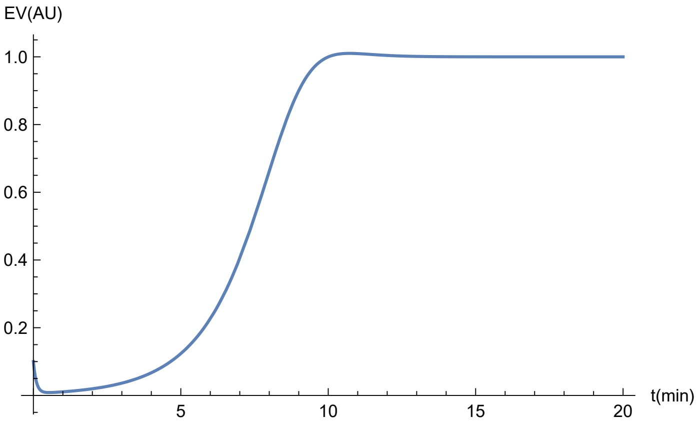


Out[48]=

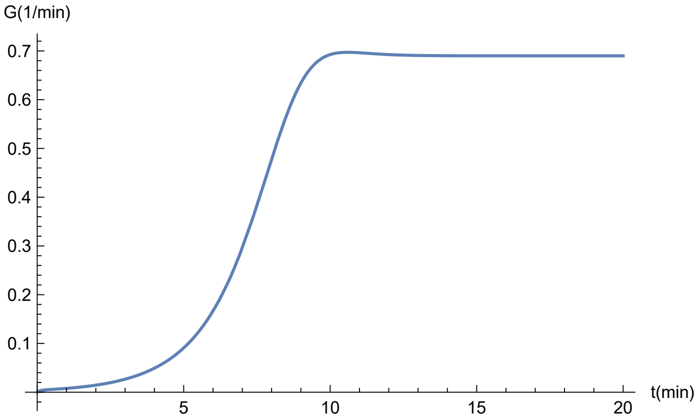


Out[49]=

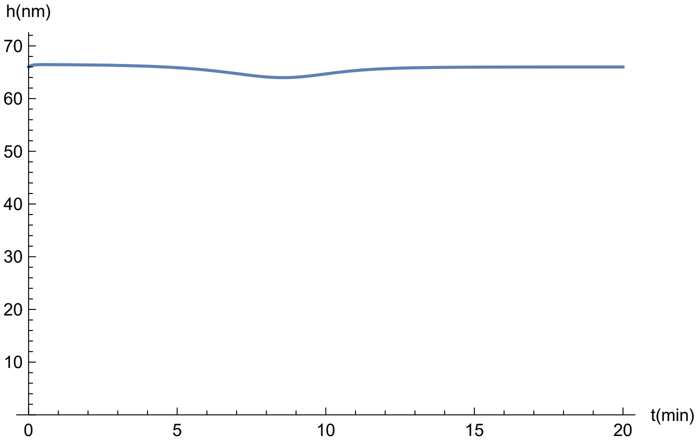

```
Export[
 "/Users/arezki/Documents/Recherche/Yeast/Louis/SimulationGraphs/SideBranching1.xlsx"
 , Table[{t, EV[t] /. sol, G[t] /. sol, 10^3 h[t] /. sol}, {t, 0, 20, 0.01}]]
```

Out[ ]= /Users/arezki/Documents/Recherche/Yeast/Louis/SimulationGraphs/SideBranching1.xlsx

We then consider initially no exocytic vesicles and smaller initial thickness to see whether this can trigger initiation of growth. The outcome is that no outgrowth occurs unless there is some level of exocytic vesicle. Very low initial level of exocytic vesicle delays the establishment of growth.

In[50]:=

```
sol = First@
  NDSolve[{EV'[t] == ϕ G[t] - α EV[t], G'[t] == If[G[t] > 0, η EV[t] (1 - θ h[t]) / h[t]^2 -
          γ (2 - θ h[t]) / (1 - θ h[t]) / h[t] G[t] × EV[t] + G[t]^2 / (1 - θ h[t]), 0],
       h'[t] == γ EV[t] - G[t] × h[t] , EV[0] == 0, G[0] == 0, h[0] == hData / 2} /.
      ConstraintsFromData /. ParametersFromData /. ϕguess, {EV, G, h}, {t, 0, 20}]
Plot[EV[t] /. sol, {t, 0, 20}, PlotRange → All, AxesLabel → {"t(min)", "EV(AU)"}]
Plot[G[t] /. sol, {t, 0, 20}, PlotRange → All, AxesLabel → {"t(min)", "G(1/min)"}]
Plot[10^3 h[t] /. sol, {t, 0, 20},
 PlotRange → {0, 1.1 × 10^3 hData /. ConstraintsFromData},
 AxesLabel → {"t(min)", "h(nm)"}]
```

Out[50]= {EV → InterpolatingFunction[ Domain: {{0., 20.}} Output: scalar ],

G → InterpolatingFunction[ Domain: {{0., 20.}} Output: scalar ],

h → InterpolatingFunction[ Domain: {{0., 20.}} Output: scalar ]}

Out[51]=

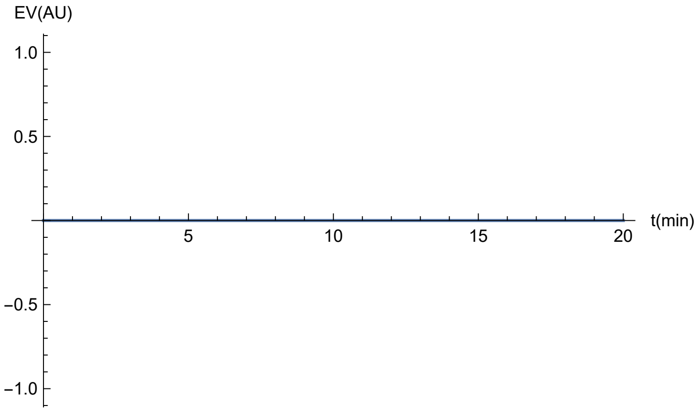

Out[52]=

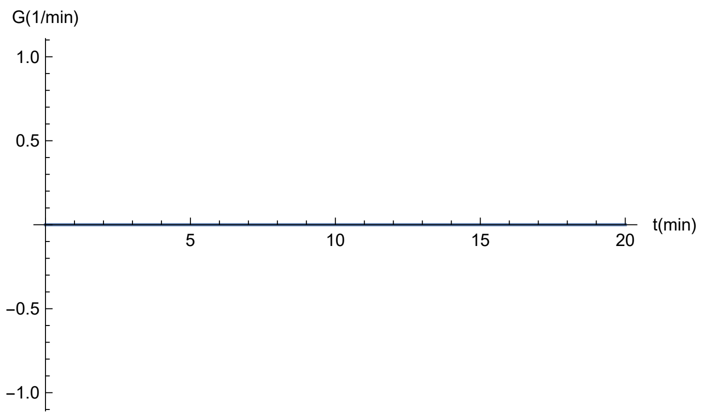


Out[53]=

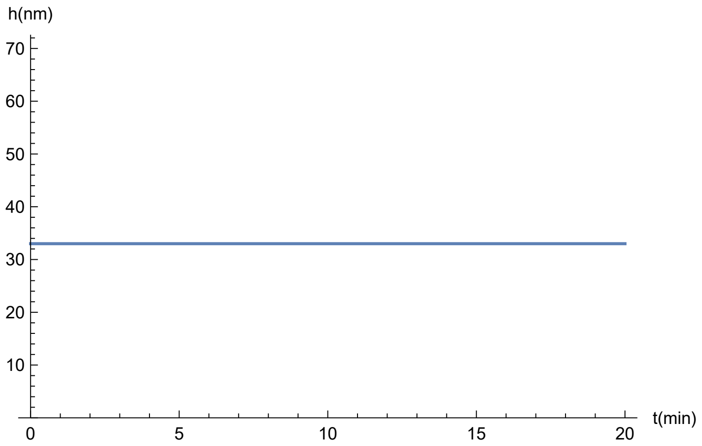


```
Export[
 "/Users/arezki/Documents/Recherche/Yeast/Louis/SimulationGraphs/SideBranching2.xlsx"
 , Table[{t, EV[t] /. sol, G[t] /. sol, 10^3 h[t] /. sol}, {t, 0, 20, 0.01}]]
```

Out[*]= /Users/arezki/Documents/Recherche/Yeast/Louis/SimulationGraphs/SideBranching2.xlsx

In[54]:=

```
sol = First@
   NDSolve[{EV'[t] == ϕ G[t] - α EV[t], G'[t] == If[G[t] > 0, η EV[t] (1 - θ h[t]) / h[t]^2 -
            γ (2 - θ h[t]) / (1 - θ h[t]) / h[t] G[t] × EV[t] + G[t]^2 / (1 - θ h[t]), 0],
        h'[t] == γ EV[t] - G[t] × h[t] , EV[0] == 0.001, G[0] == 0, h[0] == hData} /.
       ConstraintsFromData /. ParametersFromData /. ϕguess, {EV, G, h}, {t, 0, 20}]
Plot[EV[t] /. sol, {t, 0, 20}, PlotRange → All, AxesLabel → {"t(min)", "EV(AU)"}]
Plot[G[t] /. sol, {t, 0, 20}, PlotRange → All, AxesLabel → {"t(min)", "G(1/min)"}]
Plot[10^3 h[t] /. sol, {t, 0, 20},
 PlotRange → {0, 1.1 × 10^3 hData /. ConstraintsFromData},
 AxesLabel → {"t(min)", "h(nm)"}]
```

Out[54]= {EV → InterpolatingFunction[ Domain: {{0., 20.}} Output: scalar ],

G → InterpolatingFunction[ Domain: {{0., 20.}} Output: scalar ],

h → InterpolatingFunction[ Domain: {{0., 20.}} Output: scalar ]}

Out[55]=

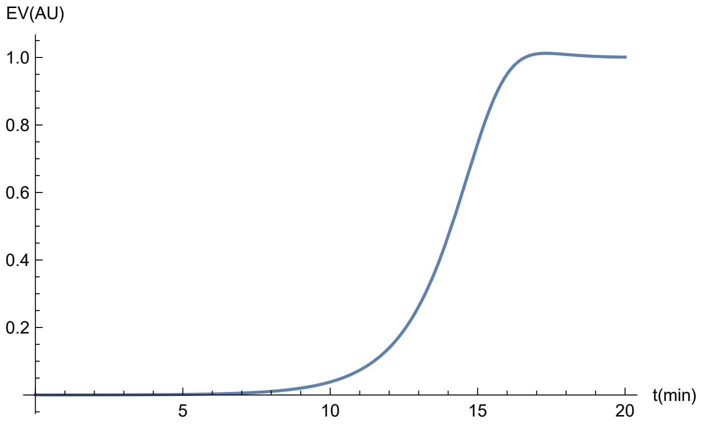


Out[56]=

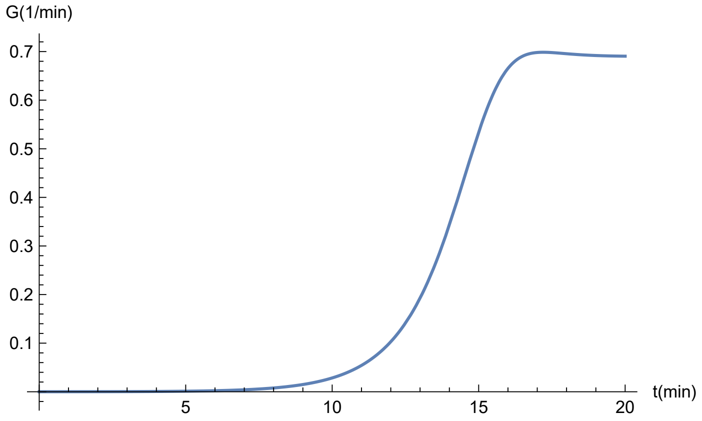


Out[57]=

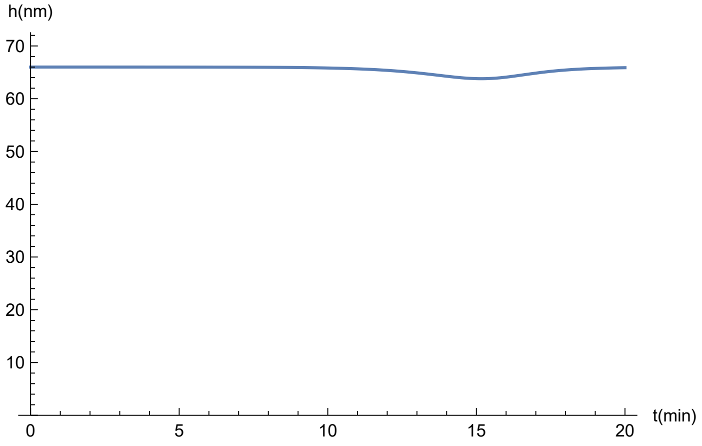

```
Export[
 "/Users/arezki/Documents/Recherche/Yeast/Louis/SimulationGraphs/SideBranching3.xlsx"
 , Table[{t, EV[t] /. sol, G[t] /. sol, 10^3 h[t] /. sol}, {t, 0, 20, 0.01}]]
```

Out[ ]= /Users/arezki/Documents/Recherche/Yeast/Louis/SimulationGraphs/SideBranching3.xlsx

### Simulation of benomyl treatment / SarA mutant

We take the reference equilibrium for the initial values. We consider that both conditions reduce the feeding and the recycling of the exocytic vesicles. At time *t*=5 min, we decrease the values of the two corresponding parameters, $\phi$ and $\alpha$, by one order of magnitude. We found that growth is slowed down only if the shift of $\alpha$ is greater than for $\phi$ (for instance -90% for $\phi$ and -91% for $\alpha$, or -60% and -70%). There is an initial peak for *EV*, which would be too small to be observed experimentally. *EV* and *G* decay to 0 (or converge to non-zero values for smaller shifts). The magnitude of increase in thickness is in the same range as in experiments and depends on the values of the shifts.

Function that changes from 0 to x at t=5

In[58]:=
```
ff[t_, x_] := 1 - (1 - x) HeavisideTheta[t - 2]
Plot[ff[t, .5], {t, 0, 20}, PlotRange → {0, 1.1}]
```

Out[59]=

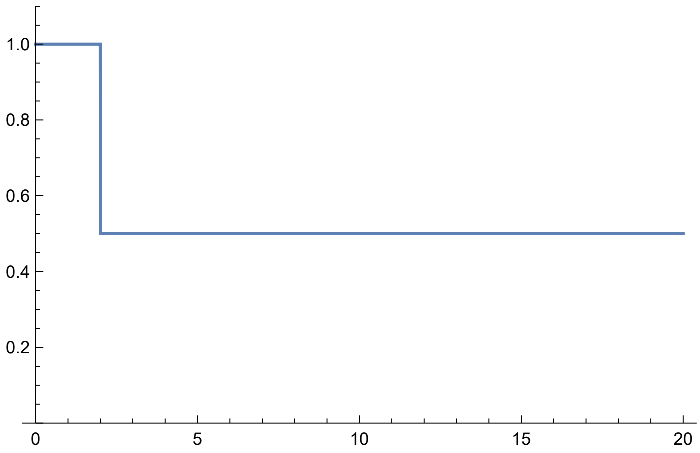


In[60]:=
```
sol = First@
  NDSolve[{EV'[t] == ϕ G[t] - α EV[t], G'[t] == If[G[t] > 0, η EV[t] (1 - θ h[t]) / h[t]^2 -
          γ (2 - θ h[t]) / (1 - θ h[t]) / h[t] G[t] × EV[t] + G[t]^2 / (1 - θ h[t]), 0],
        h'[t] == γ EV[t] - G[t] × h[t] , EV[0] == EVData, G[0] == GData, h[0] == hData} /.
       {ϕ → ϕ ff[t, .2], α → α ff[t, .19]} /. ConstraintsFromData /.
     ParametersFromData /. ϕguess, {EV, G, h}, {t, 0, 20}]
Plot[EV[t] /. sol, {t, 0, 20}, PlotRange → {0, 1.5}, AxesLabel → {"t(min)", "EV(AU)"}]
Plot[G[t] /. sol, {t, 0, 20}, PlotRange → All, AxesLabel → {"t(min)", "G(1/min)"}]
Plot[10^3 h[t] /. sol, {t, 0, 20}, PlotRange → {0, 200}, AxesLabel → {"t(min)", "h(nm)"}]
```

Out[60]=
```
{EV → InterpolatingFunction[ Domain: {{0., 20.}} Output: scalar ],
 G → InterpolatingFunction[ Domain: {{0., 20.}} Output: scalar ],
 h → InterpolatingFunction[ Domain: {{0., 20.}} Output: scalar ]}
```

Out[61]=
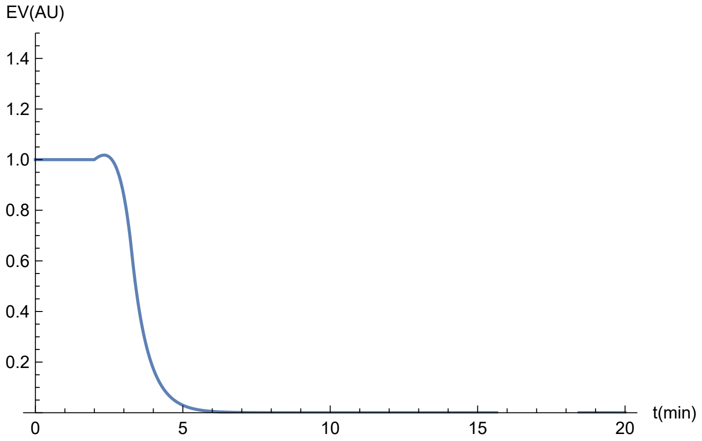


Out[62]=
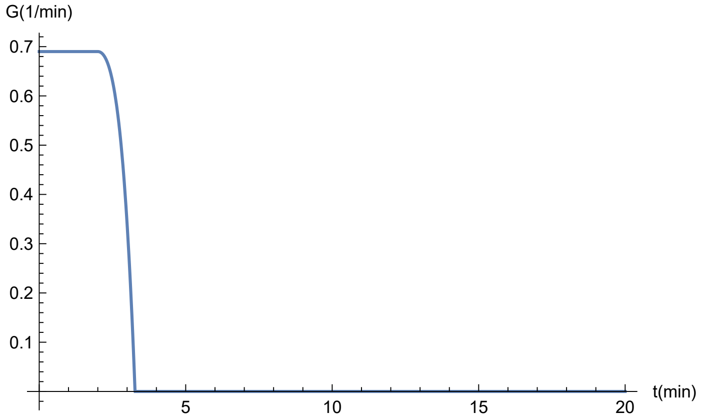


Out[63]=
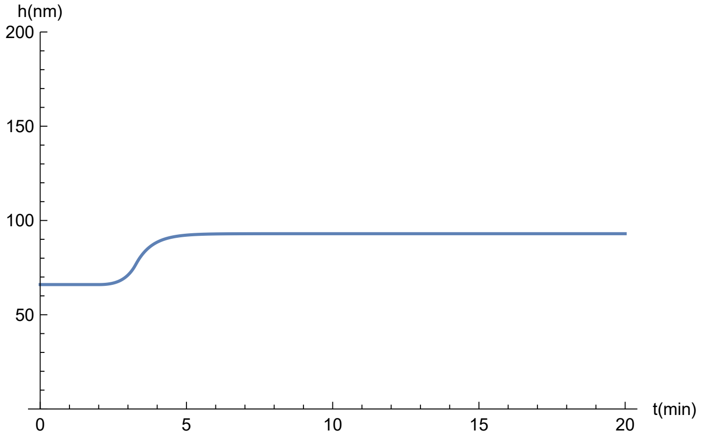


```
Export[
 "/Users/arezki/Documents/Recherche/Yeast/Louis/SimulationGraphs/BenomylSarA1.xlsx",
 Table[{t, EV[t] /. sol, G[t] /. sol, 10^3 h[t] /. sol}, {t, 0, 20, 0.01}]]
```

Out[ ]= /Users/arezki/Documents/Recherche/Yeast/Louis/SimulationGraphs/BenomylSarA1.xlsx

In[64]:=
```
sol = First@
  NDSolve[{EV'[t] == ϕ G[t] - α EV[t], G'[t] == If[G[t] > 0, η EV[t] (1 - θ h[t]) / h[t]^2 -
          γ (2 - θ h[t]) / (1 - θ h[t]) / h[t] G[t] × EV[t] + G[t]^2 / (1 - θ h[t]), 0],
        h'[t] == γ EV[t] - G[t] × h[t] , EV[0] == EVData, G[0] == GData, h[0] == hData} /.
       {ϕ → ϕff[t, .1], α → αff[t, .095]} /. ConstraintsFromData /.
     ParametersFromData /. ϕguess, {EV, G, h}, {t, 0, 80}]
Plot[EV[t] /. sol, {t, 0, 40},
 PlotRange → {-0.1, 1.5}, AxesLabel → {"t(min)", "EV(AU)"}]
Plot[G[t] /. sol, {t, 0, 40}, PlotRange → All, AxesLabel → {"t(min)", "G(1/min)"}]
Plot[10^3 h[t] /. sol, {t, 0, 40}, PlotRange → {0, 250}, AxesLabel → {"t(min)", "h(nm)"}]
```

Out[64]= {EV → InterpolatingFunction[ Domain: {{0., 80.}} Output: scalar ],

G → InterpolatingFunction[ Domain: {{0., 80.}} Output: scalar ],

h → InterpolatingFunction[ Domain: {{0., 80.}} Output: scalar ]}

Out[65]=

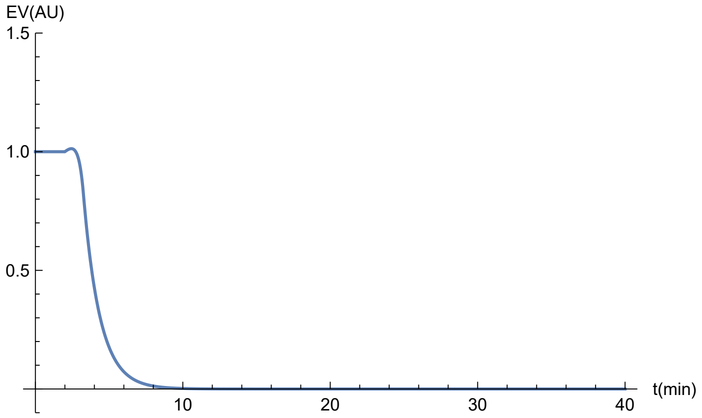


Out[66]=

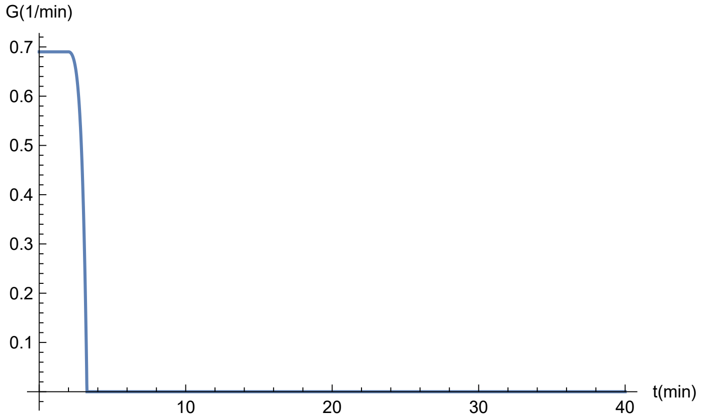

Out[67]=

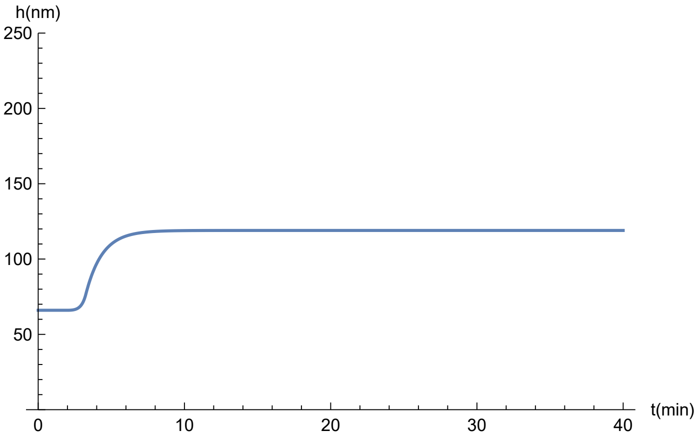


```
Export[
 "/Users/arezki/Documents/Recherche/Yeast/Louis/SimulationGraphs/BenomylSarA2.xlsx",
 Table[{t, EV[t] /. sol, G[t] /. sol, 10^3 h[t] /. sol}, {t, 0, 20, 0.01}]]
```

Out[•]= /Users/arezki/Documents/Recherche/Yeast/Louis/SimulationGraphs/BenomylSarA2.xlsx

### Simulation of osmotic shocks

We take the reference equilibrium for the initial values. In experiments, the pressure is reduced by 28%, but as this goes below the threshold for growth, we cap this decrease to 14% (which corresponds to the threshold), so that $\eta$ is multiplied by 0.86 and $\theta$ divided by 0.86 at *t*=2 min; in addition *G* is discontinuous according to Eq. (5). We find that growth stops (consistent because we used it to compute the value of $\theta$), *EV* decays to 0, and that thickness has a small increase.

In[68]:=
```
p = 0.14
sol =
 First@NDSolve[{EV'[t] == ϕ G[t] - α EV[t], G'[t] == If[G[t] > 0, -p G[t] / (1 - θ h[t])
        DiracDelta[t - 1.99999999] + η EV[t] (1 - θ h[t]) / h[t]^2 -
       γ (2 - θ h[t]) / (1 - θ h[t]) / h[t] G[t] × EV[t] + G[t]^2 / (1 - θ h[t]), 0],
     h'[t] == γ EV[t] - G[t] × h[t] , EV[0] == EVData, G[0] == GData, h[0] == hData} /.
    {η → η ff[t, 1 - p], θ → θ ff[t, 1 / (1 - p)]} /. ConstraintsFromData /.
   ParametersFromData /. ϕguess, {EV, G, h}, {t, 0, 20}]
Plot[EV[t] /. sol, {t, 0, 10}, PlotRange → All, AxesLabel → {"t(min)", "EV(AU)"}]
Plot[G[t] /. sol, {t, 0, 10}, PlotRange → All, AxesLabel → {"t(min)", "G(1/min)"}]
Plot[10^3 h[t] /. sol, {t, 0, 20}, PlotRange → {0, 100}, AxesLabel → {"t(min)", "h(nm)"}]
```

Out[68]= 0.14

Out[69]= {EV → InterpolatingFunction[ Domain: {{0., 20.}} Output: scalar ],

G → InterpolatingFunction[ Domain: {{0., 20.}} Output: scalar ],

h → InterpolatingFunction[ Domain: {{0., 20.}} Output: scalar ]}

Out[70]=

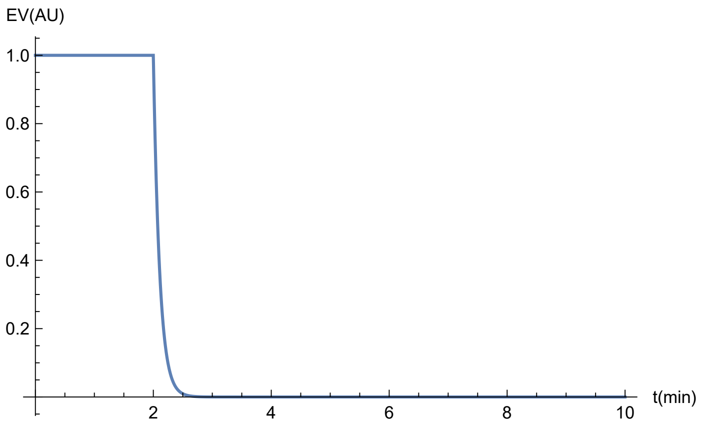


Out[71]=

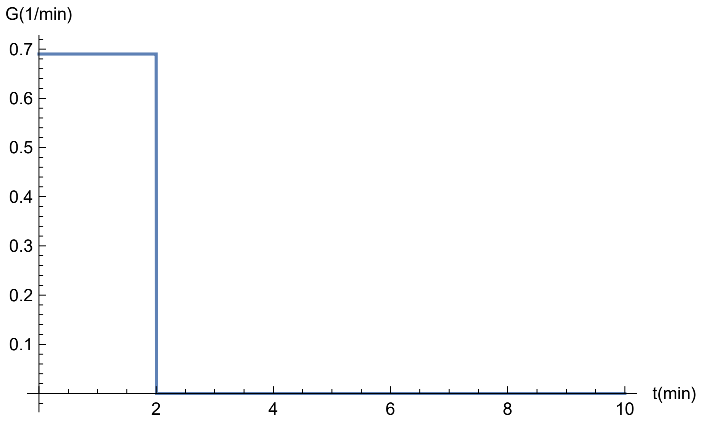


Out[72]=

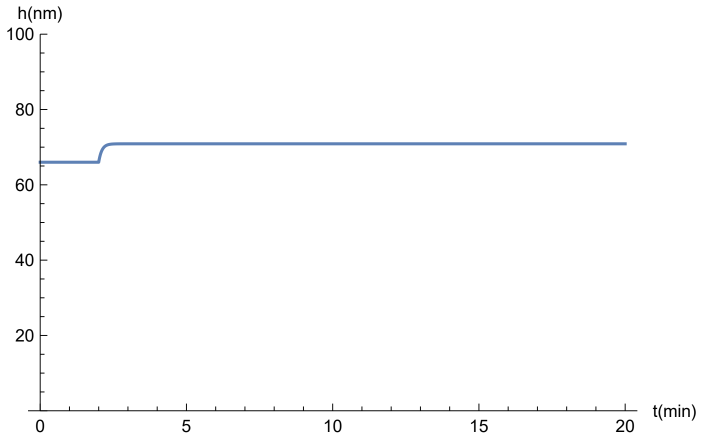

```
Export[
 "/Users/arezki/Documents/Recherche/Yeast/Louis/SimulationGraphs/OsmoticShock.xlsx",
 Table[{t, EV[t] /. sol, G[t] /. sol, 10^3 h[t] /. sol}, {t, 0, 20, 0.01}]]
```

Out[ ]= /Users/arezki/Documents/Recherche/Yeast/Louis/SimulationGraphs/OsmoticShock.xlsx

### Simulation of osmotic shocks with adaptation

We take the reference equilibrium for the initial values. We consider the same decrease in pressure as above, except that we assume that it occurs with a time scale of 6 s (to avoid numerical instabilities); in addition, we assume that pressure is restored to its initial value (by osmotic adjustment) with a timescale of 2 min. Because the state with no growth is stationary, we keep a small positive threshold for growth ($1-\theta h > 0.001$) to avoid being trapped in this no-growth state; biologically, this indirectly accounts for sources of noise. We find that growth stops and *EV* decays to 0, before recovering their initial values, and that thickness has a small transient increase.

In[73]:=
```
ff2[t_, x_] := If[t < 2, 1, x + (1 - x) Exp[-10 (t - 2)] + (1 - x) (1 - Exp[-(t - 2) / 2])]
Plot[ff2[t, .5], {t, 0, 20}, PlotRange → {0, 1.5}]
```

Out[74]=
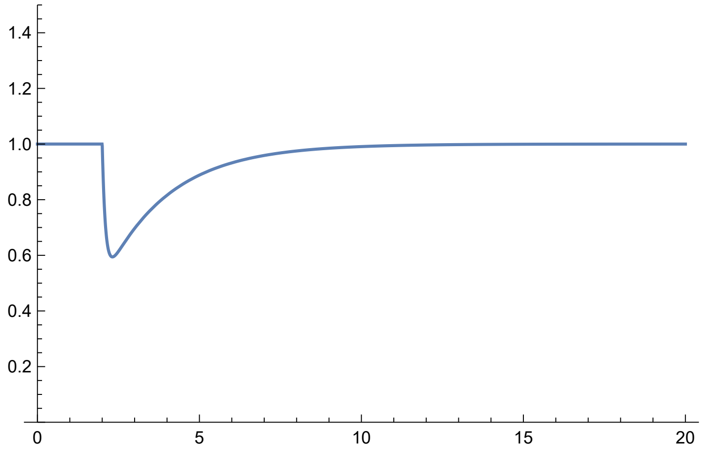


In[75]:=
```
p = 0.14
sol = First@NDSolve[
   {EV'[t] == ϕ G[t] - α EV[t], G'[t] == If[1 - θ h[t] > 0.001, η EV[t] (1 - θ h[t]) / h[t]^2 -
        γ (2 - θ h[t]) / (1 - θ h[t]) / h[t] G[t] × EV[t] + G[t]^2 / (1 - θ h[t]), 0.],
      h'[t] == γ EV[t] - G[t] × h[t] , EV[0] == EVData, G[0] == GData, h[0] == hData} /.
     {η → η ff2[t, 1 - p], θ → θ ff2[t, 1 / (1 - p)]} /. ConstraintsFromData /.
    ParametersFromData /. ϕguess, {EV, G, h}, {t, 0, 20}]
Plot[EV[t] /. sol, {t, 0, 20}, PlotRange → All, AxesLabel → {"t(min)", "EV(AU)"}]
Plot[G[t] /. sol, {t, 0, 20}, PlotRange → All, AxesLabel → {"t(min)", "G(1/min)"}]
Plot[10^3 h[t] /. sol, {t, 0, 20}, PlotRange → {0, 100}, AxesLabel → {"t(min)", "h(nm)"}]
```

Out[75]= 0.14

Out[76]= {EV → InterpolatingFunction[ Domain: {{0., 20.}} Output: scalar ],

G → InterpolatingFunction[ Domain: {{0., 20.}} Output: scalar ],

h → InterpolatingFunction[ Domain: {{0., 20.}} Output: scalar ]}

Out[77]=

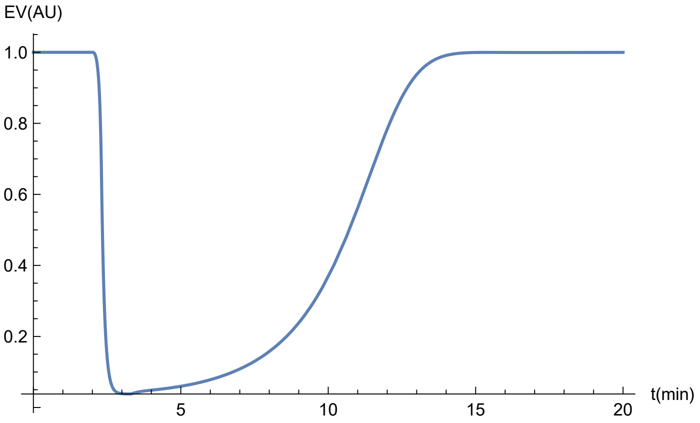


Out[78]=

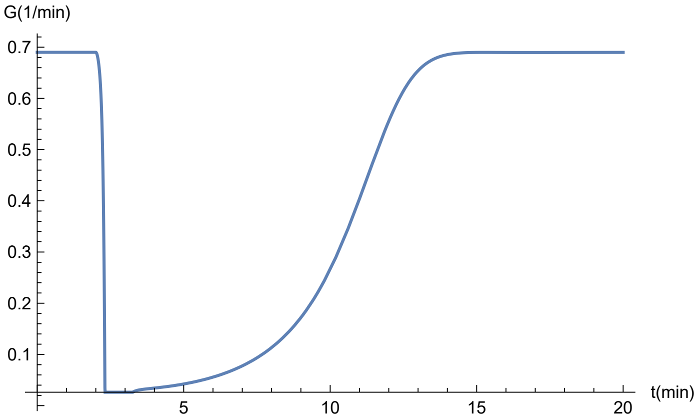


Out[79]=

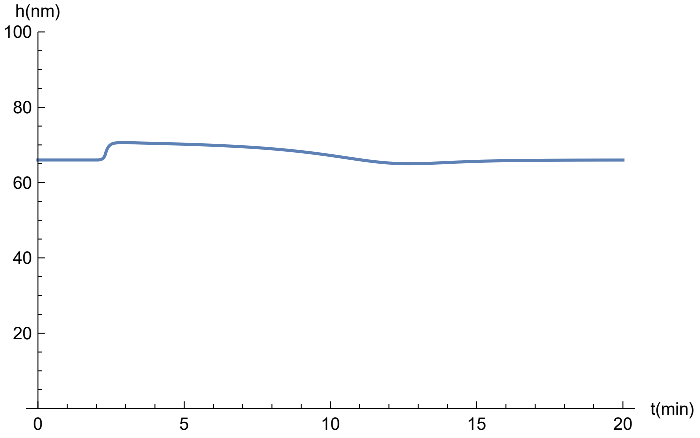

```
Export["/Users/arezki/Documents/Recherche/Yeast/Louis/SimulationGraphs/
    OsmoticShockAdaptation.xlsx",
  Table[{t, EV[t] /. sol, G[t] /. sol, 10^3 h[t] /. sol}, {t, 0, 20, 0.01}]]
```

Out[◦]= /Users/arezki/Documents/Recherche/Yeast/Louis/SimulationGraphs/OsmoticShockAdaptation .xlsx

### Estimation of parameters for germling tubes

We follow the same approch as for the reference conditions, assuming that the yield strain $\epsilon$ remains unchanged.

1) We take:
Stationary exocytic vesicle level EV = 0.14,
Stationary growth rate G = Gs/R =0.14 $\mu$m/min/0.92$\mu$m=0.15min $^{-1}$,
Stationary tip thickness h = 0.073 $\mu$m,
From Eq. (4) $\theta$ is proportional to $Y/PR$, so is multiplied by 0.73: $\theta =$ 13.03x0.73=9.51$\mu$m $^{-1}$
We then obtain values of $\gamma$, $\eta$ and a relation between $\alpha$ and $\phi$.

2) We assume the two last eigenvalues to be equal and obtain the value of $\phi$.

In[92]:=
```
ConstraintsFromDataGT = {EVData → .14, GData → 0.15, hData → 0.073, θ → 9.51}
Solve[{EVSta == EVData, GSta == GData, hSta == hData}, {γ, η, α}] // Simplify
ParametersFromDataGT = % /. ConstraintsFromDataGT // Flatten
EigenvaluesWithParametersGT =
  EigenvaluesNoGrowth /. ParametersFromDataGT /. {θ → 9.51} // Simplify
Solve[EigenvaluesWithParametersGT⟦2⟧ == EigenvaluesWithParametersGT⟦3⟧, ϕ]
ϕguessGT = %⟦2, 1⟧
```

Out[92]= {EVData → 0.14, GData → 0.15, hData → 0.073, θ → 9.51}

Out[93]= $\left\{\left\{\gamma \to \frac{\text{GData hData}}{\text{EVData}},\ \eta \to \frac{\text{GData}^2\ \text{hData}^2}{\text{EVData} - \text{EVData hData}\ \theta},\ \alpha \to \frac{\text{GData}\ \phi}{\text{EVData}}\right\}\right\}$

Out[94]= {γ → 0.0782143, η → 0.00280095, α → 1.07143 ϕ}

Out[95]= $\left\{-0.15,\ 0.170282 - 0.535714\,\phi - 6.39269\sqrt{0.000709532 - 0.00839708\,\phi + 0.00702261\,\phi^2},\ 0.170282 - 0.535714\,\phi + 6.39269\sqrt{0.000709532 - 0.00839708\,\phi + 0.00702261\,\phi^2}\right\}$

Out[96]= {{ϕ → 0.0914991}, {ϕ → 1.10422}}

Out[97]= ϕ → 1.10422

In[98]:=
```
Append[ParametersFromData /. ϕguess, ϕguess]
Append[ParametersFromDataGT /. ϕguessGT, ϕguessGT]
```

Out[98]= {γ → 0.04554, η → 0.0148114, α → 9.30581, ϕ → 13.4867}

Out[99]= {γ → 0.0782143, η → 0.00280095, α → 1.18309, ϕ → 1.10422}

In germling tubes, we find an increase by a factor of about 2 for $\gamma$, and reductions by factors of about 8 for $\alpha$ and 12 for $\phi$. Given Eq. (4), the reduction by 5.3 of $\eta$, together with the measured increase by 1.35 of *P/Y*, implies a reduction by a factor of about 7 of the product $\beta\,\mu$. In short, reduced rates (factors in the range 3-12), except for an increase in synthesis at the membrane, may explain all measured differences between germling tubes and hyphae.

### Estimation of parameters for ΔMyoV

We follow the same approch as for the reference conditions, assuming that the yield strain $\epsilon$ remains unchanged.

1) We take:
Stationary exocytic vesicle level $EV = 0.23$,
Stationary growth rate $G = Gs/R$ =0.32 $\mu$m/min/1.6$\mu$m=0.19 min $^{-1}$,
Stationary tip thickness $h = 0.081$ $\mu$m,
From Eq. (4) $\theta$ is proportional to $Y/PR$, so is multiplied by 0.68: $\theta =$ 13.03x0.68=8.86$\mu$m $^{-1}$
We then obtain values of $\gamma$, $\eta$ and a relation between $\alpha$ and $\phi$.

2) We assume the two last eigenvalues to be equal and obtain the value of $\phi$.

```
In[100]:= ConstraintsFromDataMyoV = {EVData → .23, GData → 0.19, hData → 0.081, θ → 8.86}
Solve[{EVSta == EVData, GSta == GData, hSta == hData}, {γ, η, α}] // Simplify
ParametersFromDataMyoV = % /. ConstraintsFromDataMyoV // Flatten
EigenvaluesWithParametersMyoV =
  EigenvaluesNoGrowth /. ParametersFromDataMyoV /. {θ → 8.86} // Simplify
Solve[EigenvaluesWithParametersMyoV[[2]] == EigenvaluesWithParametersMyoV[[3]], ϕ]
ϕguessMyoV = %[[2, 1]]
```

Out[100]= {EVData → 0.23, GData → 0.19, hData → 0.081, θ → 8.86}

Out[101]= $\left\{\left\{\gamma \to \frac{\text{GData hData}}{\text{EVData}}, \eta \to \frac{\text{GData}^2\,\text{hData}^2}{\text{EVData} - \text{EVData hData}\,\theta}, \alpha \to \frac{\text{GData}\,\phi}{\text{EVData}}\right\}\right\}$

Out[102]= $\{\gamma \to 0.066913, \eta \to 0.00364735, \alpha \to 0.826087\,\phi\}$

Out[103]= $\left\{-0.19, 0.241474 - 0.413043\,\phi - 7.47238\sqrt{0.00104429 - 0.00638354\,\phi + 0.00305544\,\phi^2}, 0.241474 - 0.413043\,\phi + 7.47238\sqrt{0.00104429 - 0.00638354\,\phi + 0.00305544\,\phi^2}\right\}$

Out[104]= $\{\{\phi \to 0.178912\}, \{\phi \to 1.91033\}\}$

Out[105]= $\phi \to 1.91033$

```
In[106]:= Append[ParametersFromData /. ϕguess, ϕguess]
Append[ParametersFromDataMyoV /. ϕguessMyoV, ϕguessMyoV]
```

Out[106]= $\{\gamma \to 0.04554, \eta \to 0.0148114, \alpha \to 9.30581, \phi \to 13.4867\}$

Out[107]= $\{\gamma \to 0.066913, \eta \to 0.00364735, \alpha \to 1.5781, \phi \to 1.91033\}$

In ΔMyoV, we find a small increase for $\gamma$, a reduction by factors in the range 6-7 for $\alpha$ and for $\phi$. Given Eq. (4), the reduction by 4.07 of $\eta$, together with the measured decrease by 0.95 of *P/Y*, implies a reduction by a factor of about 4 of the product $\beta\,\mu$. In short, reduced trafficking level (similar reduction by a factor in the range 4-7 of all coefficients related to trafficking) in MyoV may explain all measured differences with wild-type. Smaller in $\gamma$ suggests that cell wall synthesis is not significantly affected in this mutant background.
